# Transcriptional profiling and single-cell chimerism analysis identifies human tissue resident T cells in the human skin after allogeneic stem cell transplantation

**DOI:** 10.1101/2020.04.11.037101

**Authors:** Gustavo P. de Almeida, Peter Lichtner, Sophia Mädler, Chang-Feng Chu, Christina E. Zielinski

## Abstract

Tissue resident memory T cells (T_RM_) have recently emerged as crucial cellular players for host defense in a wide variety of tissues and barrier sites. Mouse models revealed that they are maintained long-term *in loco* unlike recirculating effector memory T cells (T_EM_). Insights into the maintenance and regulatory checkpoints of human tissue resident T cells (T_RM_) remain scarce, especially due to the obstacles in tracking T cells over time and system-wide in humans. We present a clinical model that allowed us to overcome these limitations. We demonstrate that allogeneic stem cell transplantation resulted in compartmentalization of host T cells in the human skin despite complete donor T cell chimerism in the blood, thus unmasking long-term persistence of tissue resident T cell subsets of host origin within the diverse skin T cell community. Single-cell transcriptional profiling paired with single-cell chimerism analysis provided an in-depth characterization of these *bona fide* skin resident T cells. Their phenotype, functions and regulatory checkpoints may serve therapeutic strategies for the treatment of autoimmune diseases and chronic infections, where their specific depletion versus maintenance, respectively, will have to be harnessed pharmaceutically.

## Introduction

Allogeneic hematopoietic stem cell transplantation (allo-HSCT) is a therapeutic strategy to cure various hematologic malignancies ^1^. More than 1.4 million transplants have been performed so far worldwide ^2^. Myeloablative chemotherapy, often paired with total body irradiation, precedes this procedure in order to allow for replacement of the host immune system by a donor immune system that reconstitutes from its stem cells. Engraftment of donor stem cells is routinely monitored by chimerism analysis of the blood and bone marrow, which also enables monitoring of disease relapse (i.e. leukemia). Genotyping of short tandem repeats (STRs) or microsatellites, repeat units of 2 to 7 bp in length, is commonly used to detect and quantify chimerism, due to the highly polymorphic nature that allows for easy distinction of the DNA from different individuals.

Although most fundamental insights into the properties and regulation of the human immune system stem from investigations of blood, only a small proportion of immune cells is indeed recirculating. It is therefore questionable whether the blood and bone marrow compartment provide a representative snapshot of the human immune system. In fact, only 2 % of all human T cells are present in the blood, whereas the vast majority of T cells is sequestered in peripheral tissues where they may reside long-term as resident memory T cells (T_RM_) ^3^. Landmark studies with parabiosis experiments in mouse models have provided compelling evidence for the long-term persistence of tissue resident T cells in a quiescent state ^4,5^. The obstacles in tracking T cells over time and system-wide in humans have, however, precluded *bona fide* evidence for long-*term in situ* lodgement of T_RM_ cells, in particular in the skin, and hampered insights into their homeostasis and regulatory checkpoints.

Recently, enhanced expression of ATP-binding cassette (ABC) family multidrug transporters has been observed in T_RM_. They efflux proteins and small molecules, and thus maintain cellular homeostasis as well as increased resistance to chemotherapies ^5,6^. This prompts the question whether T_RM_ might evade chemotherapy induced depletion. Likewise, it challenges the current concept that complete donor T cell chimerism, which is routinely observed upon myeloablative therapy and allo-HSCT ^7^, is representative of the overall immune system and therefore indicative of the assumed complete reconstitution of the immune system with donor stem cells.

Immunomonitoring of immune cells in peripheral tissues, such as the skin, has so far not been performed in settings of allo-HSCT. Considering the possibility of T_RM_ persistence after myeloablative conditioning chemotherapy as a consequence of lymphocyte quiescence and increased expression of multidrug transporters, tissue T cell chimerism could provide *in vivo* evidence for the existence of *bona fide* T_RM_ by their chimeric host status. In this study, we demonstrate that the clinical model of allo-HSCT can be leveraged for the identification and phenotypic, functional and regulatory analysis of T_RM_ in the skin. Single-cell transcriptional profiling paired with single-cell chimerism analysis demonstrated compartmentalization of host T cells for more than two years in the human skin and allowed for their in-depth characterization. These insights are highly relevant to harness regulatory checkpoints of T_RM_ for long-term protective immunity at barrier tissues such as the skin and to specifically target their pathogenic memory in settings of organ restricted autoimmunity.

## Results

### Physiological composition of blood and skin immune niches after allo-HSCT

In this study we investigated a female patient more than two years (796 days) after myeloablative chemotherapy and allogeneic hematopoietic stem cell transplantation with clinically established complete donor chimerism of the blood and bone marrow immune compartments. The patient did not have any clinical or histologic signs of graft-versus-host disease (GvHD) and did not receive any immunosuppressive drugs for more than 2 years **(Table 1)**. We hypothesized that this clinical setting would allow us to assess the existence of long-term *bona fide* tissue resident T cells in the skin niche based on their chimeric host status paired with an in-depth characterization of their phenotype, function and regulation.

**Table 1.**

| **Patient ID** | **Nr. 08** |
| --- | --- |
| Patient age | 38 |
| Patient sex | female |
| Donor sex | male |
| Time after allo-HSCT | 796 days |
| Diagnosis | AML |
| Induction therapy | cytarabine (7 days), daunorubicin (3 days), mitoxantrone (2 days) |
| Conditioning therapy | busulfan (4 days), cyclophosphamide (2 days), anti-thymocyte-globulin (3 days) |
| Irradiation | no |
| Initial immunosuppression | ciclosporin A, methotrexate (3 days) |
| HSCT | 8.14 x 10<sup>6</sup> CD34+ cells per kg KG |
| Donor origin | HLA-identical unrelated donor |
| HLA host | HLA-A 0201 0301<br>HLA-B 0702 4001<br>HLA-C 0304 0702<br>HLA-DRB1 0901 1501<br>HLA-DQB1 0303 0603 |
| HLA donor | HLA-A 0201 0301<br>HLA-B 0702 4001<br>HLA-C 0304 0702<br>HLA-DRB1 0901 1501<br>HLA-DQB1 0303 0603 |
| GvHD | no |
| Chimerism blood | 100% donor |
| Chimerism bone marrow | 100% donor |
| Chimerism skin | 49% host |
| CMV-Status (donor/host) | negative for host or donor prior to transplantation |
| CMV-Reactivation | negative |
| Other diseases | Vitamin D deficiency |
| Medication | Vitamin D supplementation (Dekristol) |

We first investigated the immune cell composition of the skin and blood niche of the patient in comparison to a healthy female control person. To this end, we isolated CD45^+^ cells of hematopoietic origin from the skin and blood of both donors by flow cytometry assisted cell sorting and investigated the entire 3’ single-cell transcriptomes of lymphocytes using the 10x Genomics scRNA-Seq platform **(Figure S1)**. Cell type annotation was performed by marker gene expression after Louvain clustering **(Figure S2)**. We restricted our analysis to the major lymphocyte lineages such as CD4, CD8, Treg and γδ T cells, as well as NK, NKT and B cells. Skin and blood of the allo-HSCT patient demonstrated a composition of lymphocyte subsets that was comparable to the healthy control in both quality and quantity **(Figure 1a, b)**. To validate cell type annotation based on single-cell transcriptomes, we also assessed the differential expression of protein surface markers that define immune cell lineages on skin and blood lymphocytes with a CyTOF dataset of 4-5 independent healthy donors **(Figure S3)**. This demonstrated, despite potential effects of the skin digestion procedure on protein surface marker expression, that the cell types and their respective frequencies, as identified by scRNA-seq in the patient and healthy control, were within the range of normal interindividual variation. This suggests that a physiological composition of both immune niches was evident at the time of analysis despite myeloablative chemotherapy and allo-HSCT.

**Figure 1.**
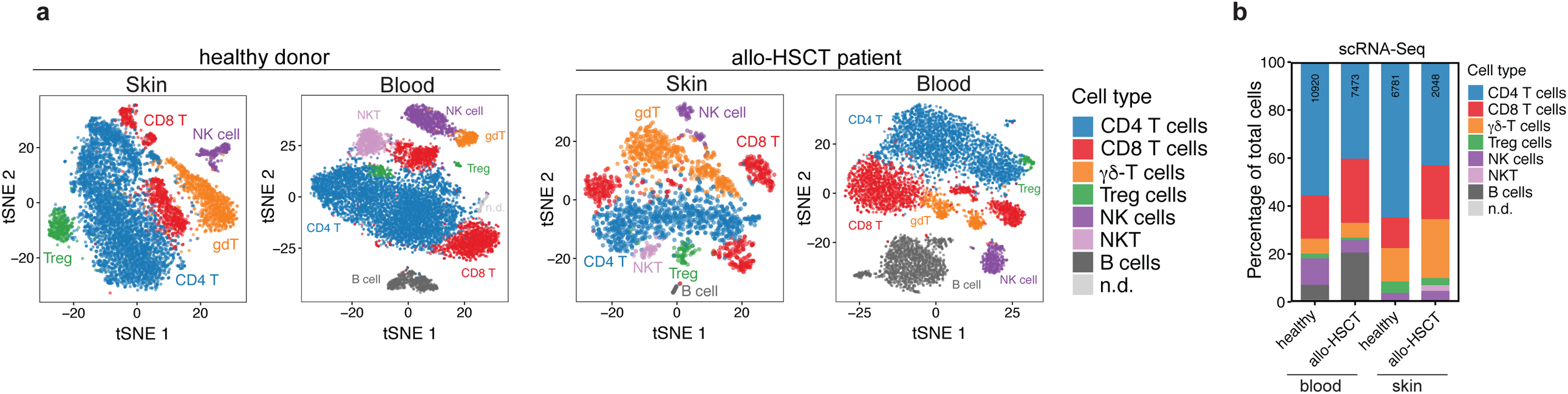
Distribution of lymphocytes subsets in the skin and blood with and without allo-HSCT. **(a)** t-SNE depicting the cluster cell identity by different color codes based on manual annotation with known marker genes as shown in (Figure S2). scRNA-seq of fresh CD45^+^ lymphocytes was performed 796 days after allo-HSCT. **(b)** Differential distribution of lymphocyte subsets in the skin and blood of the allo-HSCT patient and a healthy control person. n.d., not determined.

### Persistence of host lymphocytes in the skin after allo-HSCT unmasks skin T_RM_

After having excluded any abrogation in the composition of the immune cell niche in skin and blood in the allo-HSCT patient, we set out to prove the existence of *bona fide* skin resident human T cells (skin T_RM_). We hypothesized that the putative persistence of host lymphocytes in the human skin paired with complete donor lymphocyte chimerism in the circulating blood would demonstrate the fixed compartmentalization of T cells in the skin and thus their long-term skin T cell residency (> 2 years) **(Figure 2a)**. We first isolated CD3^+^ T cells from the blood and skin of the patient (706 and 1096 days after allo-HSCT) and tested for T cell chimerism by short-tandem-repeat-PCR using 16 markers, which allows for robust distinction of DNA from different sources due to the highly polymorphic nature of the STRs (microsatellites) **(Table 2a, b)**. Skin fibroblasts cultured from the CD45^−^ fraction of the punch biopsy served as a positive control for host DNA. 14 out of 16 markers provided detectable signals that allowed us to calculate total donor T cell chimerism **(Figure 2a, b)**. In accordance with the common dogma that complete donor T cell chimerism follows myeloablative chemotherapy and allo-HSCT upon successful immune reconstitution, we detected 100 % donor CD3^+^ T cells in the blood and bone marrow (not shown). Interestingly, we observed that only 43% of skin CD3^+^ T cells were of donor origin. This demonstrated that a high proportion of host CD3^+^ T cells were maintained in the skin compartment without recirculation. 10 months later (1096 after allo-HSCT), we obtained 51% donor chimerism in support of a rather stable maintenance of the host skin T cell population over time **(Table 2b)**. This provided evidence for the existence of a *bona fide* skin resident T cell subset (skinT_RM_).

**Table 2.** a 709 days after allo-HSCT

| Marker | Donor Chimerism (%) |
| --- | --- |
| AMEL | 39 |
| CSF1PO | 44 |
| D13S317 | 32 |
| D16S539 | - |
| D18S51 | 52 |
| D19S433 | 48 |
| D21S11 | 45 |
| D2S1338 | 47 |
| D3S1358 | 39 |
| D5S818 | - |
| D7S820 | 49 |
| D8S1179 | 42 |
| FGA | 44 |
| TH01 | 42 |
| TPOX | 35 |
| vWA | 40 |
| Median | 43 |

**b** 1096 days after allo-HSCT
| Marker | Donor Chimerism (%) |
| --- | --- |
| AMEL | 51 |
| CSF1PO | 57 |
| D13S317 | 48 |
| D16S539 | - |
| D18S51 | 58 |
| D19S433 | 56 |
| D21S11 | 50 |
| D2S1338 | 58 |
| D3S1358 | 49 |
| D5S818 | - |
| D7S820 | 52 |
| D8S1179 | 51 |
| FGA | 53 |
| TH01 | 50 |
| TPOX | 47 |
| vWA | 46 |
| Median | 51 |

**Figure 2.**
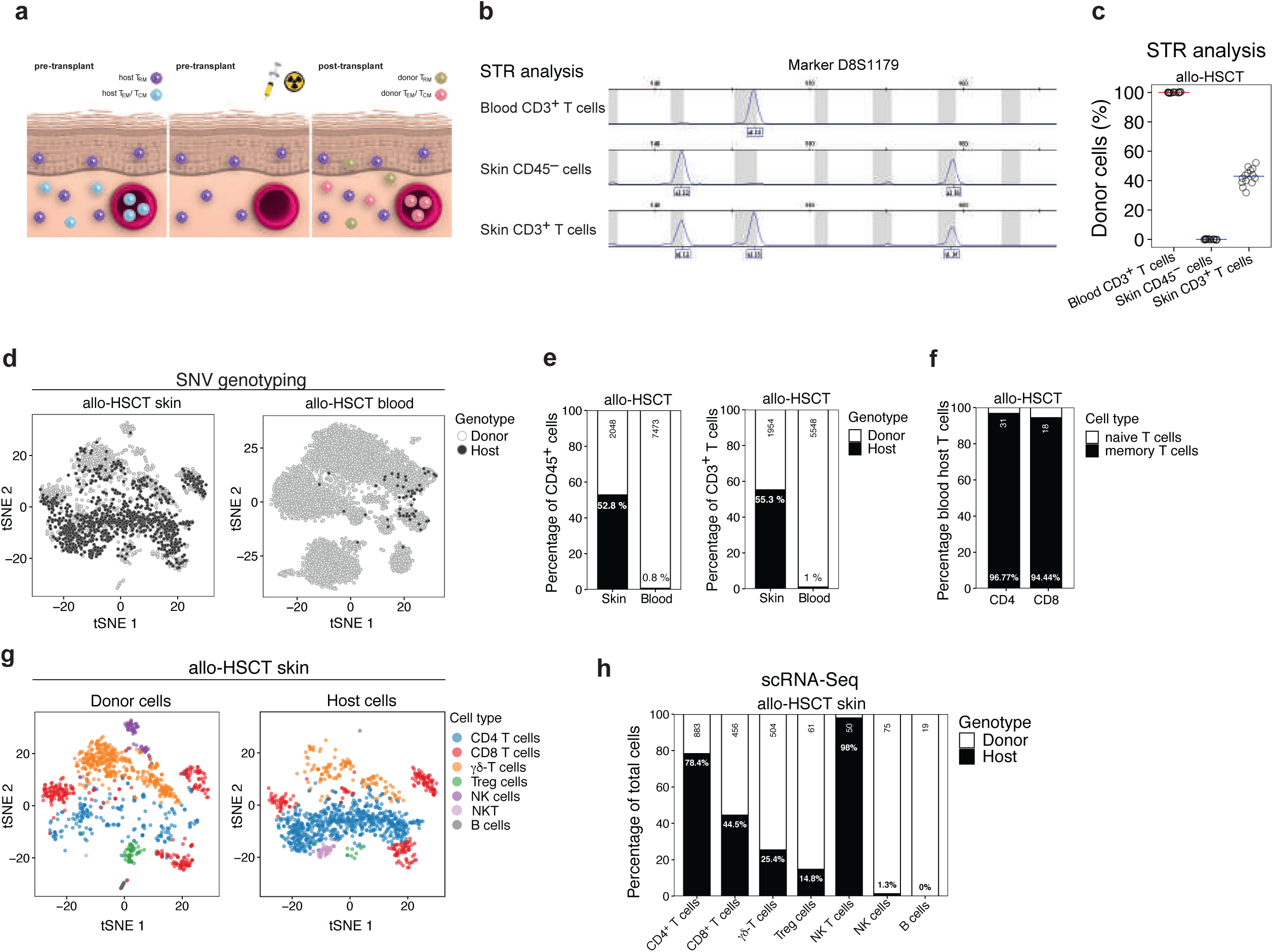
Chimerism analysis of lymphocytes identifies host cells as resident cells. **(a)** Clinical model of allo-HSCT. Skin punch biopsy for lymphocyte isolation and chimerism analysis was performed 706 days after allo-HSCT. **(b)** Polymerase chain reaction of short tandem repeats (STR-PCR) of CD3^+^ T cells from the blood and skin and of CD45^−^ cells from the skin. Shown is the electropherogram of one representative locus out of 16 analyzed loci. **(c)** Cumulative analysis of 14 distinct detectable microsatellite loci. Shown is the median percentage of donor cells of total cells calculated from peak heights and peak areas of each of the analyzed 14 loci. **(d)** Single-cell genotyping through SNV using snpclust 796 days after allo-HSCT. **(e)** Distribution of host and donor cells identified through SNV in single-cell transcriptomes of lymphocytes from blood and skin in the allo-HSCT patient. The number inside the bar indicates the percentage of host cells within total lymphocytes (left) or total T cells (right), while the number on top indicates total lymphocytes (left) or total T cells (right) analyzed. **(f)** Distribution of naïve versus memory T cells within the CD4^+^ and CD8^+^ T cell lineage within host blood cells as identified by the average cluster expression of subset-specific genes. **(g)** Cell identity annotation was performed by average cluster expression of cell-type specific markers within host versus donor lymphocytes, which was visualized in the reduced dimensional space calculated with tSNE. **(h)** Genotype distribution within each of the annotated cell subsets in the skin. The number inside the black bar indicates the percentage of host cells within total cells of the respective cell subset. The number on top indicated the total cell number within the respective cell subsets.

To corroborate our findings, we performed single-cell RNA sequencing (scRNA-seq) 796 days after allo-HSCT, which allowed us to interrogate the host versus donor status of each individual lymphocyte isolated from skin or blood based on the detection of small nuclear variants (SNV). 52.8% of all skin lymphocytes were of host origin, while almost all blood lymphocytes (99.2 %) were of donor origin, in line with the information provided by STR-PCR analysis **(Figure 2c, d)**. This relative distribution of host versus donor cells also applied to the CD3^+^ T cell compartment (55.3 % host T cells in skin) **(Figure 2d)**. Interestingly, the scRNA-seq method was able to unmask the existence of a small population of 1% host T cells within circulating blood T cells **(Figure 2c, d)**. It distributed equally between the CD4 and CD8 T cell lineage and displayed a memory differentiation state **(Figure 2e)**.

The single-cell transcriptomes allowed further annotation of lymphocyte subsets, which segregated into host and donor cells. This analysis revealed a differential distribution of these lymphocyte subsets within the host or donor cell compartments **(Figure 2f, g)**. Interestingly, we observed a high fraction of host CD4^+^ T cells (78.4 %) compared to a relatively low fraction of host CD8^+^ T cells (44.5 %), suggesting preferential tissue maintenance of the T helper cell lineage as compared to the cytotoxic CD8^+^ T cell lineage. Surprisingly, host Treg cells only had a minor representation within the total skin Treg population (14.8 %). The NK and B cell compartment displayed only very few cells of host origin. NKT cells, in contrast, were strongly skewed towards the host genotype (98 %). However, due to their low absolute cell numbers in the skin, this fraction of host NKT cells has to be interpreted with caution.

We next asked whether sex-mismatched allo-HSCT also allows to identify resident lymphocytes by sex-associated genes. Donor stem cells were derived from a male donor. We classified individual lymphocytes as male after interrogation of 17 out of 90 Y-linked genes, which were detectible in at least one cell of the blood of the allo-HSCT patient **(Figure 3a, b)**. This demonstrated a high overlap in SNV and sex-gene based analysis for the identification of host lymphocytes. Interestingly, the Y-linked gene RPS4Y1 was sufficient to robustly identify male donor lymphocytes **(Figure 3c, d)**. Likewise, XIST, which is exclusively expressed in female cells for inactivation of one X-chromosome, was able to identify female host cells **(Figure 3e, f)**. This demonstrates that sex-mismatched transplantation settings allow the identification of host and thus resident lymphocytes with high fidelity by only targeted RPS4Y1 or XIST gene transcript detection.

**Figure 3.**
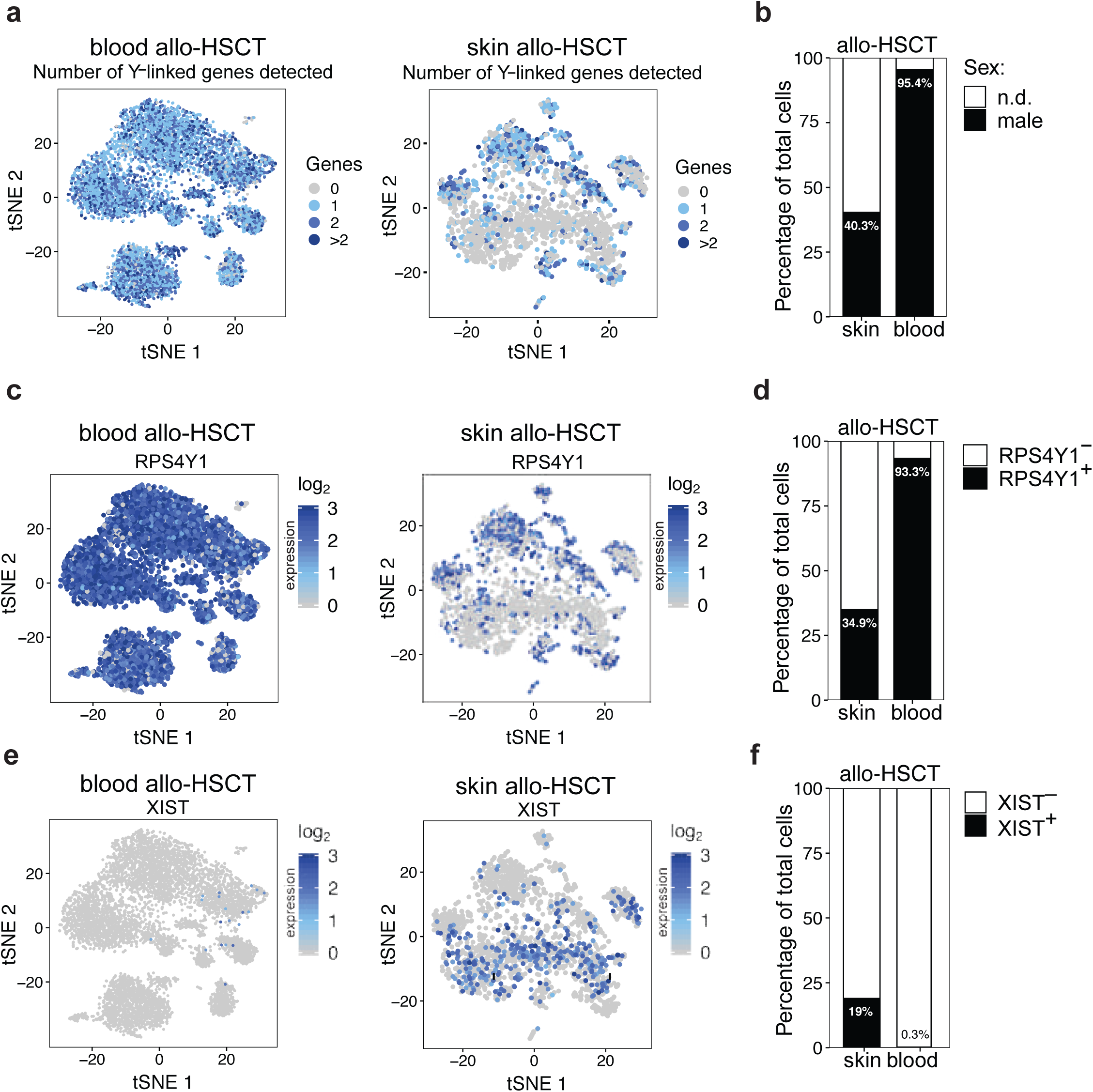
Sex-gene associated transcripts can identify resident skin lymphocytes in sex-mismatched transplantation. **(a)** Total number of 17 Y-linked genes from the HUGO database (Gene Nomenclature Committee at the European Bioinformatics Institute) with at least one read count detected in each cell from blood (left) and skin (right) of the allo-HSCT patient shown after dimensionality reduction of scRNAseq data by tSNE. 90 out of 501 protein-coding and non-coding Y-linked genes were present in the annotation for the reference assembly used for read mapping. 17 out of these 90 genes were detected in at least one cell from the blood of the allo-HSCT patient and then used for sex-genotyping. **(b)** Distribution of cells expressing at least one Y-linked gene (male) in the scRNA-Seq data or no detectable Y-linked genes (n.d.). The number inside the bar indicates the percentage of male cells our of total cells. **(c)** Cpm-normalized single-cell expression of the Y-linked gene RPS4Y1 after dimensionality reduction with tSNE. **(d)** Distribution of cells according to the expression status of the Y-linked gene RPS4Y1 in the scRNA-Seq data. The number inside the bar indicates the percentage of RPS4Y1-expressing cells within total analyzed lymphocytes. **(e, f)** Lymphocytes were analyzed for gene expression of XIST according to **(c, d)**.

Overall, these data demonstrate by multiple analysis strategies the existence of *bona fide* tissue resident lymphocytes in the human skin and suggest a differential replacement kinetics by distinct donor lymphocyte subsets.

### Host T cells in the skin and blood after allo-HSCT enrich for T cell residency signatures

After having identified skin resident lymphocyte populations by their long-term sequestration in the skin, we interrogated them on the single-cell level for transcriptomic core residency versus core recirculation signatures. The core signatures have previously been established by the identification of common overlapping genes that are consistently up- or downregulated (residency or circulation, respectively) in T cells from human tissues compared to secondary lymphoid organs and blood as well as in several murine tissues as compared to circulating splenic memory T cells ^8 9,10^. According to our hypothesis that host lymphocytes represent long-term resident cells, whereas donor cells constitute a mixture of recirculating as well as newly established resident lymphocytes, we detected a significant enrichment of the core residency signature in the host lymphocyte population **(Figure 4a, c, Figure S4a, b)**. Likewise, the core circulatory signature was decreased in host as compared to donor cells. This pattern was consistent among CD4, CD8 and γδ T cell subsets **(Figure 4b)**. We did, however, not observe an enrichment for the core residency signature in Treg cells.

**Figure 4.**
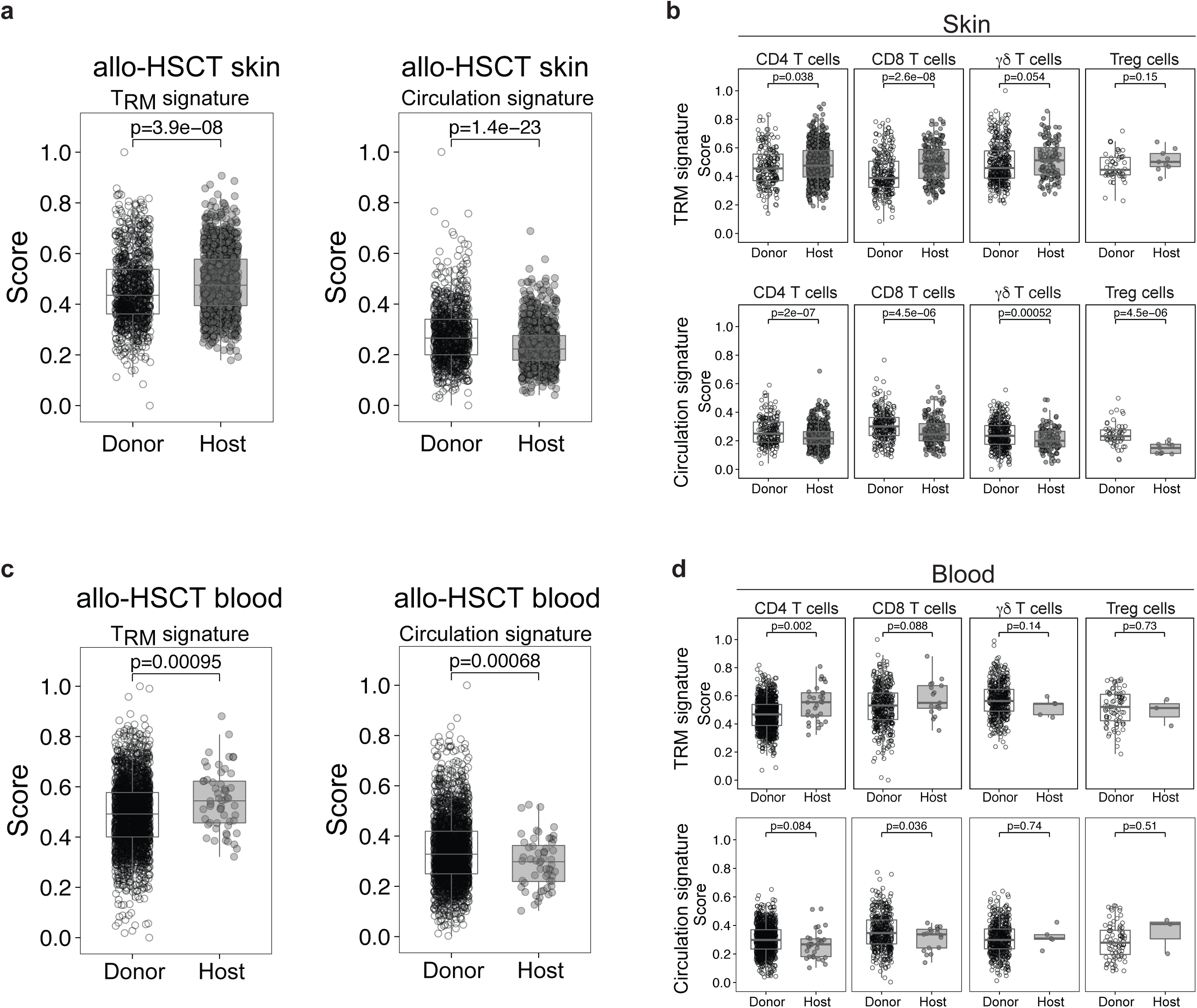
Host lymphocytes in skin and blood after allo-HSCT enrich for T cell residency signatures. **(a, c)** Single-cell gene expression scores for the core residency signature (left) and the core circulation signature (right) for all T cells identified by marker gene expression based annotation after Louvain clustering of the scRNA-seq data from the skin of the allo-HSCT patient. The core signatures were established from T_RM_ cells from skin, lung, gut, kidney, small intestine and brain in comparison to circulating splenic memory T cells. The overlapping genes, that are upregulated in all the tissues constitute the residency signature, whereas the downregulated genes represent the circulating signature ^36^. The cells were split between donor and host cells as identified by snpclust and were compared to each other using two-tailed unpaired Student’ t-test. **(b, d)** Single-cell gene expression scores for the core residency signature and core circulation signature for individual T cell subsets identified by marker gene expression based annotation after Louvain clustering.

Interestingly, we also observed that the core residency signature was significantly enriched in the few host lymphocytes that we could detect in the blood **(Figure 4c)**. This applied to CD4^+^ and CD8^+^ T cells, but not γδ T cells and Treg cells **(Figure 4d)**. This suggests that T_RM_ might retain some recirculation potential and that they preserve their transcriptomic identity despite displacement into the circulation. We then examined their tissue source by the expression of tissue specific signatures on the single-cell level. Despite the limitations that very low cell numbers impose (53 cells), our interrogation of lung, gut and skin tissue signatures revealed a mixed pattern for tissue identities and therefore suggested that the host T cells, which were identified in the blood by their SNV, could have been displaced from various organs harboring tissue resident T cells **(Figure S5)**. In sum, these data support a tissue residency versus circulatory program for host versus donor T cells, respectively, and suggest that resident T cells can exit into the circulation while maintaining their transcriptomic identity.

### Cell cycle analysis reveals enhanced proliferation of CD8^+^ T cells of host origin in the blood circulation

The identification of circulating host T cells in the blood with features of tissue resident T cells prompted further investigations into their potential mode of tissue exit. We reasoned that antigen specific stimulation might have initiated their mobilization into the blood allowing for further expansion and redistribution into peripheral tissues for broader spatial coverage of protective immunity. This hypothesis was triggered by a concurrent report that T_RM_ cells were detectable during a short time window after vaccination of immune donors, implicating their contribution to systemic secondary immune responses ^11^. We therefore decided to analyze the cell cycle stages of blood and skin lymphocytes of host and donor origin in our allo-HSCT patient, respectively. We observed a comparable distribution of the G0, S, and G2/M phases between the healthy donor and allo-HSCT patient 796 days after transplantation **(Figure 5a)**, which was largely dominated by resting lymphocytes in both blood and skin. No differences in cell cycle stages were notable by the comparison of host and donor lymphocytes in the skin and blood of the allo-HSCT patient **(Figure 5b)**. Interestingly, we observed an increased fraction of G2/M CD8^+^ T cells of host origin in the blood of the allo-HSCT patient **(Figure 5c)**. This population, which we have categorized above as exT_RM_ cells based on their host status and enrichment for residency signatures, therefore displayed enhanced proliferation. This was not the case for its CD4^+^ host T cell counterpart. In the healthy donor, no difference in the distribution of cell cycle stages was notable between CD4^+^ and CD8^+^ T cells in the skin and blood **(Figure 5c)**. Taken together, these data demonstrate that putative exT_RM_ cells of the CD8^+^ T cell lineage can expand in the circulation based on single-cell genomic cell cycle analysis. As no apparent antigenic trigger was identified in our patient (i.e. negative CMV status), TCR clonality analyses will have to be conducted to determine whether antigen specific reactivation or cytokine driven polyclonal proliferation might have caused tissue exit and whether CD4^+^ T cells might follow the same rules if exposed to appropriate stimuli.

**Figure 5.**
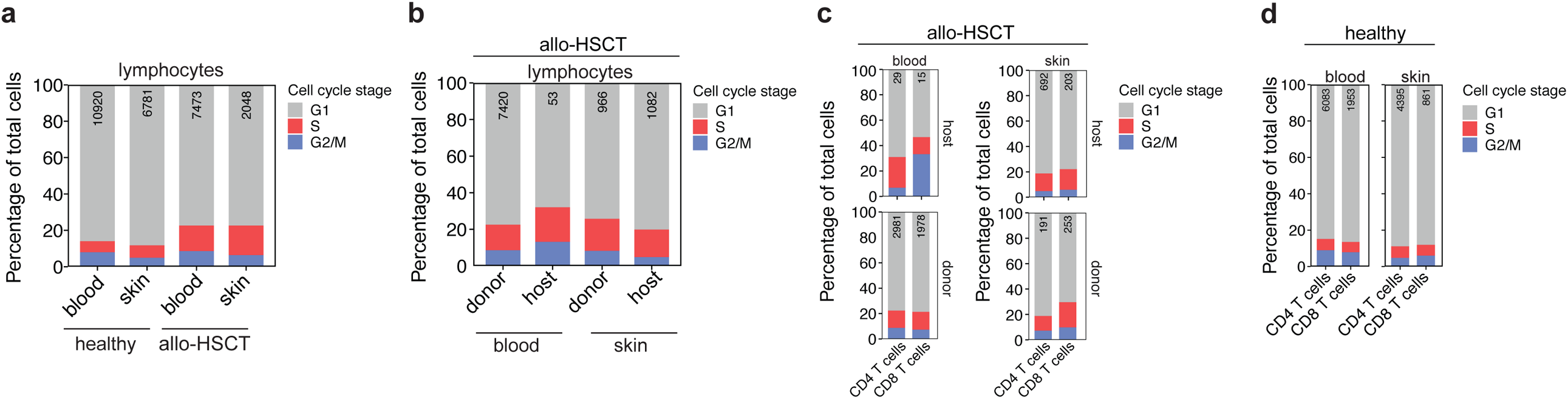
Cell cycle analysis of blood and skin lymphocytes of host and donor origin. **(a-d)** Distribution of cell cycle stages in lymphocytes and T cell subsets from blood and skin of the allo-HSCT patient and healthy donor as identified in the scRNA-Seq data. To assess the cell cycle stage at the single cell level, the Cyclone function implemented in the scran R package was used with the built-in trained gene-pairs^40^. The number on top indicate the total amount of cells detected in each sample. Host and donor cells were identified based on SNV. T cell subsets were identified by marker gene expression based annotation after Louvain clustering.

### Increased *LGALS3* expression identifies long-term resident skin lymphocytes

After having identified that a significant proportion of lymphocytes persisted in the human skin for at least two years, we aimed to dissect their identity and regulation. The possibility to distinguish theses long-term resident lymphocytes by their direct comparison to donor lymphocytes from the identical human skin microenvironment is ideal to overcome the caveats of cell harvesting from distinct tissues. In particular, previous attempts to compare tissue cells to those circulating in blood and secondary lymphoid tissues were severely affected by the impact that tissue digestive procedures impose on transcriptomes ^12^. In order to identify specific genes that correlated with skin T cell residency, we therefore analyzed differential gene expression between host and donor CD3^+^ T cells **(Figure 6a)**. As expected, the sex-associated genes *XIST* and *RPS4Y1* were differentially expressed in host versus donor T cells, respectively. Interestingly, *LGALS3* was significantly upregulated in host T cells of both of the CD4^+^ and CD8^+^ lineage **(Figure 6a-c)**. *LGALS3* encodes the protein Galectin-3, a member of the beta-galactoside-binding protein family, which plays an important role in cell-cell adhesion and cell-matrix interaction ^13^. We validated the expression of Galectin-3 on the protein level and observed a significant increase in T cells from healthy skin as compared to healthy matched blood T cells **(Figure 6d)**. This property could therefore potentially confer superior tissue maintenance. Among the top eight significantly upregulated genes within host T cells, we could also identify *PRDM1*, which encodes Blimp1, a previously reported central regulator of T cell tissue residency ^14^, as well as *IL7R*, which suggests enhanced susceptibility to IL-7 signaling and thus T cell survival ^15^. Other significantly upregulated genes that we identified by differential gene expression between host and donor T cells were *ANKRD28, RORA, SNHG7* and *KLF6*. They have so far not been assigned any roles in the regulation of skin resident T cells. Genes that were significantly downregulated in host T cells were *MTRNR2L1, GNLY, CD7, TRGC2, XCL1 and LITAF*. The selective analysis of T cells from the CD4^+^ and CD8^+^ lineage revealed increased *VIM* expression, encoding the structural protein vimentin, in host compared to donor cells. Vimentin provides resilience to cells when under mechanical stress *in vivo* and can act as an organizer of a number of critical proteins involved in cell attachment, migration and signaling ^16^. Interestingly, we found *KLRG1* to be downregulated by host CD8 T cells in the skin. Killer cell lectin-like receptor subfamily G member 1 (KLRG1) is an inhibitory receptor limiting effector functions and a marker for highly differentiated cells and end stage cells ^17^. It has also been shown to be a cadherin receptor, which can mediate Ca^2+^ dependent cell-cell adhesion ^18^.

**Figure 6.**
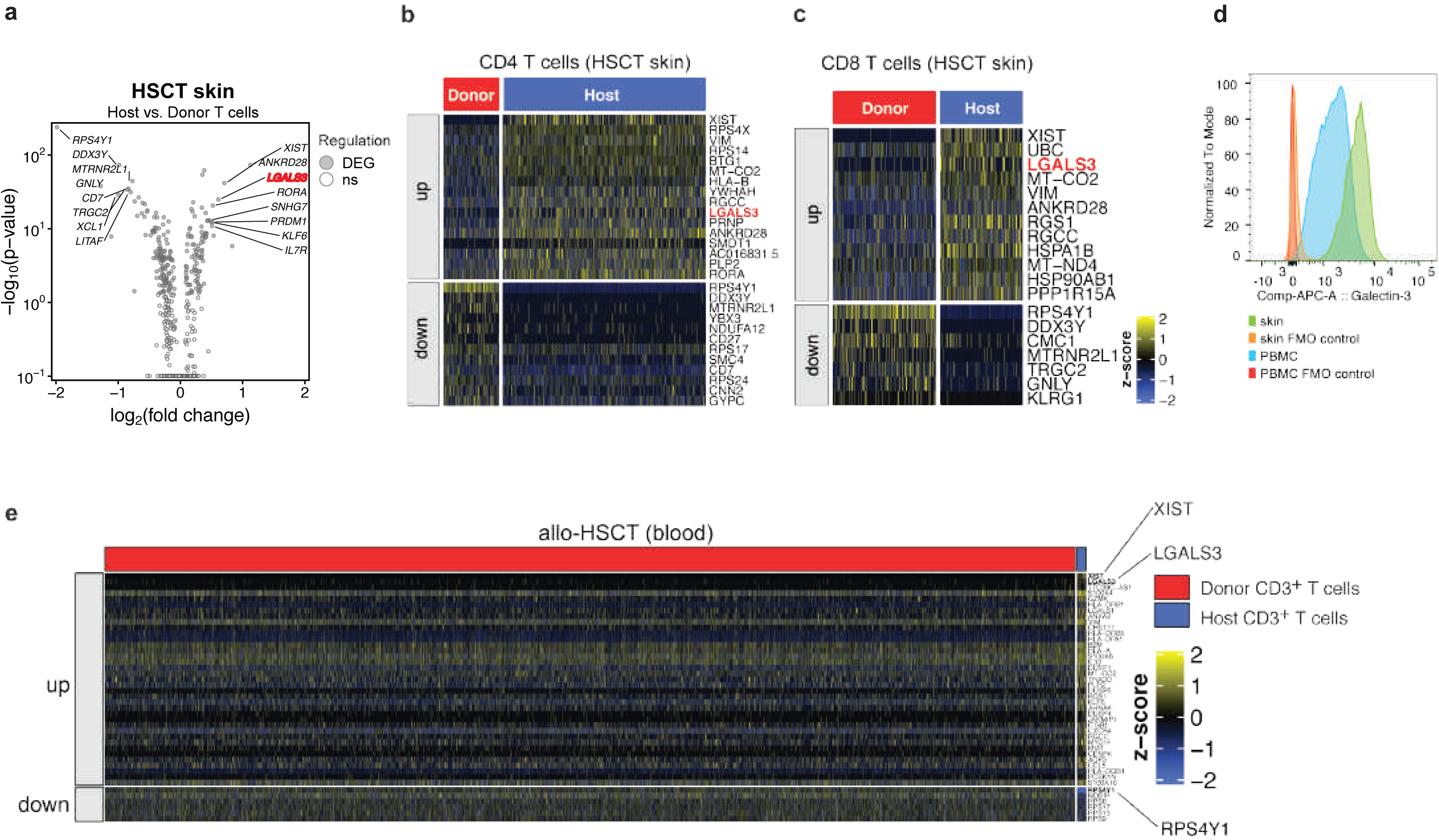
Differential gene expression between host and donor T cells in the skin niche after allo-HSCT. **(a)** Volcano plot depicting differentially expressed genes (DEG) between host and donor cells within all T cells in the skin of the allo-HSCT patient. Differentially expressed genes were identified using Wilcoxon rank-sum test implemented in the Seurat function FindAllMarkers. P-values were adjusted for multiple tests using Bonferroni correction. Genes with an adjusted p-value < 0.05 were considered significant. Shown are the top 8 significantly up- and downregulated genes. **(b, c)** Heatmap showing the scaled normalized expression values for all differentially expressed genes (adjusted p-value < 0.05) detected in host compared to donor CD4^+^ **(b)** and CD8^+^ T cells **(c)** that were identified in the skin of the allo-HSCT patient. The genes are ordered decreasingly according to their level of significance. **(d)** Flow cytometry of CD3^+^ gated viable matched PBMC and skin lymphocytes. Shown is one representative staining out of three. **(e)** Differential gene expression as in (b,c) performed between host and donor CD3^+^ T cells from the blood of the allo-HSCT patient.

Interestingly, *LGALS3* was not only enriched in host skin T cells but also in the small population of host T cells (1%), which we detected in the blood **(Figure 6e)**. The retention of this putative T_RM_ marker is therefore consistent with the overall enrichment of residency signatures and reduction in circulatory signatures within host blood T cells, thus supporting a history of tissue exit (exT_RM_) and a resident T cell lineage differentiation stage (outside-in hypothesis) ^19^.

Taken together, the transcriptome-wide comparison of host versus donor T cells on the single-cell level within the identical skin niche revealed novel regulatory candidates for the identification of *bona fide* skin resident T cells, but also confirmed targets that have previously been considered to confer residency as well as longevity to T cells in tissue niches. Considering that the donor T cell compartment of the skin comprises newly resident (< 2 years) as well circulating T cells, the overall difference in gene expression between the host and donor T cell populations was low and could unmask differences that are inherent to long-term residence.

## Discussion

Allogeneic stem cell transplantation (allo-HSCT) is potentially life-saving and curative in patients with hematological malignancies. Its full potential is limited by graft-versus-host disease (GvHD). A graft-versus-leukemia effect, however, is beneficial to clear residual disease and prevent relapses. Until now, the current paradigm suggests that myeloablative therapy and stem cell transplantation result in a replacement of the donor immune system upon successful immune reconstitution. Our data, instead, demonstrate the persistence of a large proportion of skin resident lymphocytes of host origin despite almost complete donor lymphocyte chimerism in the blood and bone marrow. This shows that a large proportion of the human immune system is sequestered as tissue resident cells from the recirculation. Considering that the human skin contains twice as many T cells as the blood compartment and that other tissues, not investigated here, could also harbor tissue resident T cells ^3^, this leaves the overall human immune system largely unaffected by allo-HSCT. The true physiological frequency and long-term maintenance of skin T_RM_ cells might have even been underestimated in our study by subclinical GvHD caused by entry of alloreactive donor T cells. Our data therefore surprisingly implicate that barrier immunity by skin T_RM_ cells is to a large extent still exerted by the host immune system.

Although insights into the regulation of human tissue resident T cells remain scarce and are mainly fueled by findings obtained in mouse models, i.e. use of CD69 and CD103 as surrogate markers of human T_RM_ ^9,20,21^, a few elegant studies have previously made the effort to track the long-term maintenance of *bona fide* human T_RM_. Donor CD8^+^ T cells were shown to persist long-term in the duodenum after pancreatic-duodenal transplantation ^22^. Likewise, donor T cells were identified in the bronchoalveolar lavage 12 months after lung transplantation supporting the long-term maintenance of lung T_RM_ ^23^. In cutaneous T cell lymphoma patients, the application of alemtuzumab, a drug that depletes most recirculating lymphocytes, revealed the persistence of CD4^+^ and CD8^+^ T cells in the unaffected skin, thus suggesting differential exit kinetics of skin T cells and even their long-term persistence ^24^. Our strategy to monitor skin T_RM_ by chimerism analysis following allo-HSCT complements these previous approaches without concurrent perturbation of the immune system with immunosuppressants or other drugs. However, the myeloablative chemotherapy that occurred more than 2 years prior to our analysis purged recirculating lymphocytes out of their tissue niches. This could have created an artificial expansion of the T_RM_ niche with abrogations in their physiological composition. Additionally, we cannot rule out that specific subpopulations within the skin T_RM_ populations might have been susceptible to the chemotherapy and are lost from our late time-point analysis.

Our results have clinical implications. Pathogenic T cell memories characterized by T cell receptor (TCR) specificities against autoantigens or allergens might likewise persist in the host T cell compartment and thus continue to exert autoimmunity or allergy until fully replaced by a healthy donor immune system. Clinical studies, however, support a cure of chronic inflammatory tissue restricted diseases such as psoriasis as well as contact allergies following allogeneic and autologous HSCT ^25,26^. Although this might challenge the concept of T_RM_ being the culprits of their pathogenesis, this phenomenon could also be explained by a higher activation state or distinct molecular architecture that predisposes autoreactive T_RM_ cells to depletion by chemotherapy or immunosuppressants.

Another intriguing clinical implication that emerges from our study is the impact of skin T_RM_ maintenance on GvHD. It has previously been shown in a cohort of 20 patients, that recipients of lung transplants experience less graft dysfunction and acute cellular rejection if they display higher frequencies of lung persisting donor T_RM_ ^23^. If analogous mechanisms are assumed between lung T_RM_ and skin T_RM_, one could speculate that a higher frequency of skin T_RM_ following allo-HSCT could protect from GvHD. Mechanistically, a higher frequency of host T cells, which has adapted to its skin niche prior to allo-HSCT, could continue to serve its homeostatic and protective functions, while limiting entry of allo-reactive and thus tissue destructive donor cells to their niche in a space and nutrient dependent manner. More patients need to undergo skin immune monitoring to provide robust correlations between maintenance of T_RM_ and GvHD. Harnessing the persistence of T_RM_ at barrier surfaces by targeting their distinct regulatory networks could therefore be a novel strategy to reduce GvHD, if this correlation holds true.

A third clinical implication refers to immunotherapies. Adoptive transfer of T cells, which are selected or engineered for their functional properties, stemness and T cell receptors, is a promising strategy for the treatment of cancers and chronic infectious diseases. Their entry into peripheral tissues, where they are expected to exert their functions, might likewise be affected by occupancy of peripheral tissue niches by T_RM_. The importance of T_RM_ for protective immunity has already been suggested half a century ago by pioneering work by McGregor and Gowans who demonstrated intact secondary immune reactions upon ablation of recirculating lymphocytes ^27^. Immunotherapies should therefore be designed to enter peripheral tissues by, first, conditioning the respective tissue niches, i.e. by creating “space”, and, second, by promoting the establishment of a tissue residency fate in the adoptively transferred T cell product.

An intriguing finding of our study was the presence of a small population of host memory T cells in the blood, which could be unmasked by the high sensitivity of our single-cell transcriptomic chimerism analysis by SNVs. Their host origin and enrichment of tissue resident gene signatures compared to donor blood T cells suggests a classification as tissue mobilized exT_RM_ cells. While the overall blood T cell population displayed a resting functional state, the host blood CD8^+^ T cell population displayed a strong enrichment of proliferating T cells according to the proportion of cells within the G2/M phase. It remains scarcely investigated whether and how T_RM_ cells exit their tissue of origin ^10,19^. Based on our cell cycle analysis, we speculate that preceding T cell activation might have mobilized them out of their respective tissues. The concomitant maintenance of tissue specific signatures suggests a potential relocation to their respective tissues of origin after proliferation. This could represent an elegant mechanism by with large tissues such as barrier organs regulate a redistribution of expanded numbers of antigen specific T cells and thus protective immunity. This would translate into more homogenous distribution of antigen specific T cells within the organ upon their reentry as well as a potential crosstalk with other organs. Interestingly, CD4^+^ T cells of host origin, which also displayed a tissue resident signature in the blood, did not display increased proliferation, suggesting either a proliferation independent mechanism of tissue exit or tissue mobilization by T cell activation in the absence of proliferation. Because we only analyzed one timepoint, previous proliferation cannot be excluded. This would be in line with a concurrent study that suggested a tight kinetics of T_RM_ tissue exit and reentry upon antigen specific T cell stimulation by vaccination ^11^.

We found a biased distribution of host versus donor cells within distinct lymphocyte lineages in the skin. CD4^+^ cells had a stronger propensity for maintenance in the skin as compared to CD8^+^ T cells. This relative distribution in favor of the CD4^+^ T cell lineage was surprising considering that particularly CD8^+^ T cells have previously been associated with residency features due to their epithelial sequestration in mice. This might, however, have been biased by their experimental generation with viral infections ^28^. Using the drug alemtuzumab, which depletes most recirculating lymphocytes in settings of cutaneous T cell lymphoma, it was has been reported that CD4^+^ T cells were much more abundant in the human skin than CD8^+^ T cells before and after depletion of the recirculating T cell pool, supporting our conclusions ^29^. Yet, CD4^+^ Treg cells, which exert anti-inflammatory functions, were to a large extend replaced by donor Treg cells. This suggests that Treg cells are less prone to establish residency. This is surprising, considering that their generation depends on TGF-β, which is known to induce the residency marker CD103 and thus increased retention and accumulation within tissues ^30^. Increased *in vivo* cycling of Treg cells as compared to memory and naïve T cells could have potentially biased their tissue elimination during myeloablative chemotherapy ^31^. The differential maintenance of other T subsets (Th1, Th2, Th17 cells) within the T helper cell lineage remains to be elucidated. We showed that NKT cells were almost exclusively of host origin, suggesting exclusion of newly entering donor NKT cells after allo-HSCT. However, the overall number of NKT cells in the skin is very low and variations in the relative proportions of host and donor T cells are possible upon examination of a larger patient cohort. B cells have previously also been reported to assume residency properties ^32^. Although we identified several B cells in the skin of the allo-HSCT patient, their scarcity precluded a reliable chimerism analysis.

Our study provided for the first time an in-depth characterization of *bona fide* skin T_RM_ cells with long-term maintenance (over 2 years) and their direct comparison to more recently lodged skin T cells and recirculating T cells within the same tissue. This unmasked *LGALS3*, encoding Galectin-3, for the first time as a novel candidate for the identification and regulation of skin T_RM_ cells. A role for Galectin-3 in the localization of T cells in tissues such as lymph nodes, bone marrow and lung has previously been suggested through its ability to confer adherence to the extracellular tissue matrix ^8^. Its *in vivo* significance for the regulation of T cell tissue residence remains to be dissected and might be facilitated in the near future by the availability of several Galectin-3 inhibitors that have already been identified and tested in animal models of cancer and various fibrotic diseases ^33^.

Cumulatively, this work exemplifies the power of exploiting a clinical situation as proof-of-concept for the existence of *bona fide* human skin T_RM_. Chimerism analysis in the context of allo-HSCT revealed the differential maintenance of lymphocyte subsets over time, tissue exit of T_RM_, and *LGALS3* as a novel gene that is significantly enriched within the skin T_RM_ population. Importantly, our results are in disagreement with the current paradigm that complete allo-HSCT replaces the host immune system in the skin and possibly other peripheral tissues. Protective as well as pathogenic immunity, which has been shaped over a lifetime, therefore continues to be provided by the host.

## Material & Methods

### Cell purification and sorting

Fresh peripheral blood was obtained from a healthy donor as well as a patient 796 days after allogeneic stem cell transplantation (allo-HSCT) following myeloablative conditioning chemotherapy with busulfan, cyclophosphamide and ATG. At the time of skin and blood sampling, the patient received Vitamin D supplementation but was not on any immunosuppressive therapy for more than a year. Peripheral blood mononuclear cells (PBMC) were isolated by density gradient centrifugation using Biocoll (Merck). Cells of hematopoietic origin were isolated as live CD45^+^ cells (clone 2D1, Biolegend) after preincubation with Human TruStain FcX (Biolegend) to a purity of over 98%. The cells were sorted with a BD FACSAria Fusion (BD Biosciences). Skin T cells from the breast and abdomen were isolated from punch biopsies following a standardized digestion protocol with the gentleMACS™ Dissociator and the whole skin dissociation kit containing enzyme A and D (both Miltenyi Biotec) before FACS-sorting for live (Hoechst 33258 negative or propidium iodide negative) CD45^+^ cells analogous to the PBMC. The gating strategy is shown in Figure S1. Galectin-3 staining was performed with human Galectin-3 Alexa-Fluor 647 conjugated antibody (R&D, clone 194801). Ethical approval was obtained from the Institutional Review Board of the Technical University of Munich (195/15s, 146/17s, 491/16s) and the Charité-Universitätsmedizin Berlin (EA1/221/11). All work involving humans was carried out in accordance with the Declaration of Helsinki for experiments involving humans.

### Single cell RNA Sequencing

CD45^+^ cells of hematopoietic origin from the skin and matched blood of both a healthy donor and a patient 796 days after allogeneic stem cell transplantation (allo-HSCT) were sorted by flow assisted cell sorting (FACS). The cells were used for library construction with the Chromium Single Cell 3’ Reagents v2 (10x Genomics, Inc). The libraries were sequenced with the Illumina system (Illumina, Inc.) according to the manufacturer’s instructions, with 150bp and 100bp of length for healthy donor samples and allo-HSCT patient samples, respectively. Read alignment and gene counting with single-cell resolution was performed with CellRanger v3.0.0 (10x Genomics, Inc), using the default parameters and the pre-built human reference v3.0.0 (10x Genomics, Inc) based on Ensembl GRCh38 release 93. The output filtered data was analyzed with the R package Seurat v3.0.0 ^34^. Inclusion of cells for further downstream analyses met the viability criteria of more than 200 and less than 2000 detected genes as well as less then 20% of mitochondrial genes within all genes. The heterogeneity associated with mitochondrial gene contamination and with the number of unique molecular identifiers (UMI) per cell were regressed out using the ScaleData function from Seurat package. The top 8000 features with the highest variance detected with the variance stabilizing transformation (vst) were used for linear dimensional reduction by PCA. The first 20 PCs were used for clustering and for non-linear dimensionality reduction by tSNE (perplexity 30). Clusters were identified by running the Louvain algorithm after constructing a KNN graph. They were annotated to specific cell types by inference based on the expression of cell type markers.

### Genotyping through single-cell RNA Sequencing

Single-cell chimerism analysis was performed by the identification and comparison of small nucleotide variants (SNV) using snpclust ^35^, with minimum quality for the SNV call set to 80 and minimum SNV base quality of 1. Only SNV detected in at least two cells were considered. The existence of two genotypes instead of only one genotype in a sample was assumed if a minimum of 90 % of the cells showed a posterior probability of at least 70 % to follow the two-genotype model. To verify whether the genotype matched that of the donor, both deduced genotypes in skin were compared to the deduced blood genotype on the basis of shared SNV.

The skin genotype with higher similarity to the blood genotype, and the cells assigned to it, were identified as donor cells, while the one with lower similarity was defined as host cells. For the sex genotyping of blood and skin samples from the allo-HSCT patient (female), who underwent a sex-mismatched transplantation, the male cells were identified according to the expression of Y-linked genes. A list of 501 protein-coding and non-coding Y-linked genes was retrieved from the HUGO Gene Nomenclature Committee (HGNC) database. A total of 90 Y-linked genes from this list were present in the annotation for the reference assembly used for read mapping. Out of those 90 genes, 17 were detected in at least one cell from the blood of the HSCT patient and, thus, were used for the sex genotyping. Cells with a minimum of one read count in at least one of the Y-linked genes were defined as male cells. For female cell identification, cells with a minimum of one read count for the gene XIST were defined as female cells.

### Gene set enrichment analysis

Single-cell gene set expression scores were calculated as the average expression of all genes from the set subtracted by the average expression of random background genes with similar range of expression values. The scores were obtained using the AddModuleScore function from Seurat package. The expression values were aggregated in 24 bins and one hundred control genes were used for each gene. The residency gene signature and circulatory gene signature were retrieved from public data sets ^8,10,36^.

### Short tandem repeat PCR (STR-PCR)

STR analysis was done with the AmpFLSTR Identifiler PCR Amplification Kit (ThermoFisher Scientific, Darmstadt, Germany) using 10 ng genomic DNA according to the manufacturer’s recommendation. This kit amplifies 15 STR markers and Amelogenin in one multiplex assay.

Amplification products were mixed with GeneScan 500 LIZ size standard (ThermoFisher Scientific, Darmstadt, Germany) and separated by size using capillary electrophoresis on an ABI3730 sequencer (ThermoFisher Scientific, Darmstadt, Germany). Data analysis was performed with GeneMapper 4.0 software (ThermoFisher Scientific, Darmstadt, Germany). This included automatic sizing and allele calling. Every electropherogram was checked manually and alleles were added or removed wherever appropriate. Peak heights and peak areas were exported for further analysis.

### CyTOF analysis

The .fcs files from the human T cell atlas ^37^ were downloaded from http://flowrepository.org (Repository ID: FR_FCM_ZZTM) and processed in R with the flowCore package ^38^. The compensated data was transformed using the flowCore “logicleTransform” function with w = 0.25, t = 16409, m = 4.5, and a = 0 as input parameters. Each of the donors was downsampled to 11930 cells using the FlowJo Plugin DownSampleV3. Non-linear dimensionality reduction by tSNE was performed with the python package scanpy ^39^ with perplexity set to 30 and the random state set to 0. For tSNE calculation the normalized protein expression values of CD19, CD3, CD4, CD8, CD45RA, CD45RO, TCRγδ, CD56 and CD25 were used as input. Cell type annotation was performed in FlowJo using the gating strategy outlined in Figure S3.

## Supplementary figures

**Figure S1.**
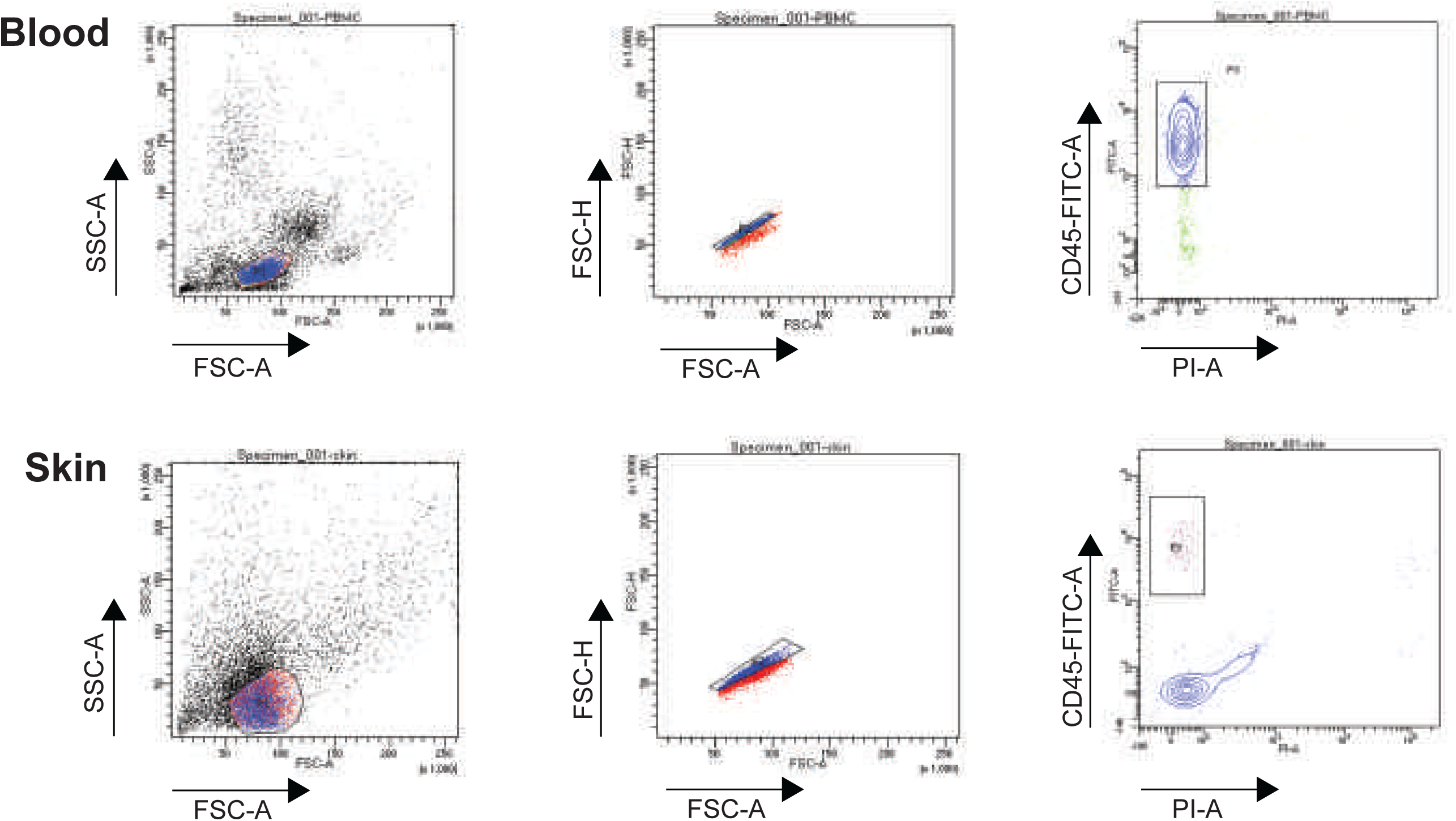
Gating strategy for isolation of lymphocytes from blood and skin. Lymphocytes were sorted as live CD45^+^ cells from skin and blood after enzymatic skin digestion or ficoll separation, respectively. The sorting strategy is representative for samples from the healthy donor and the allo-HSCT patient.

**Figure S2.**
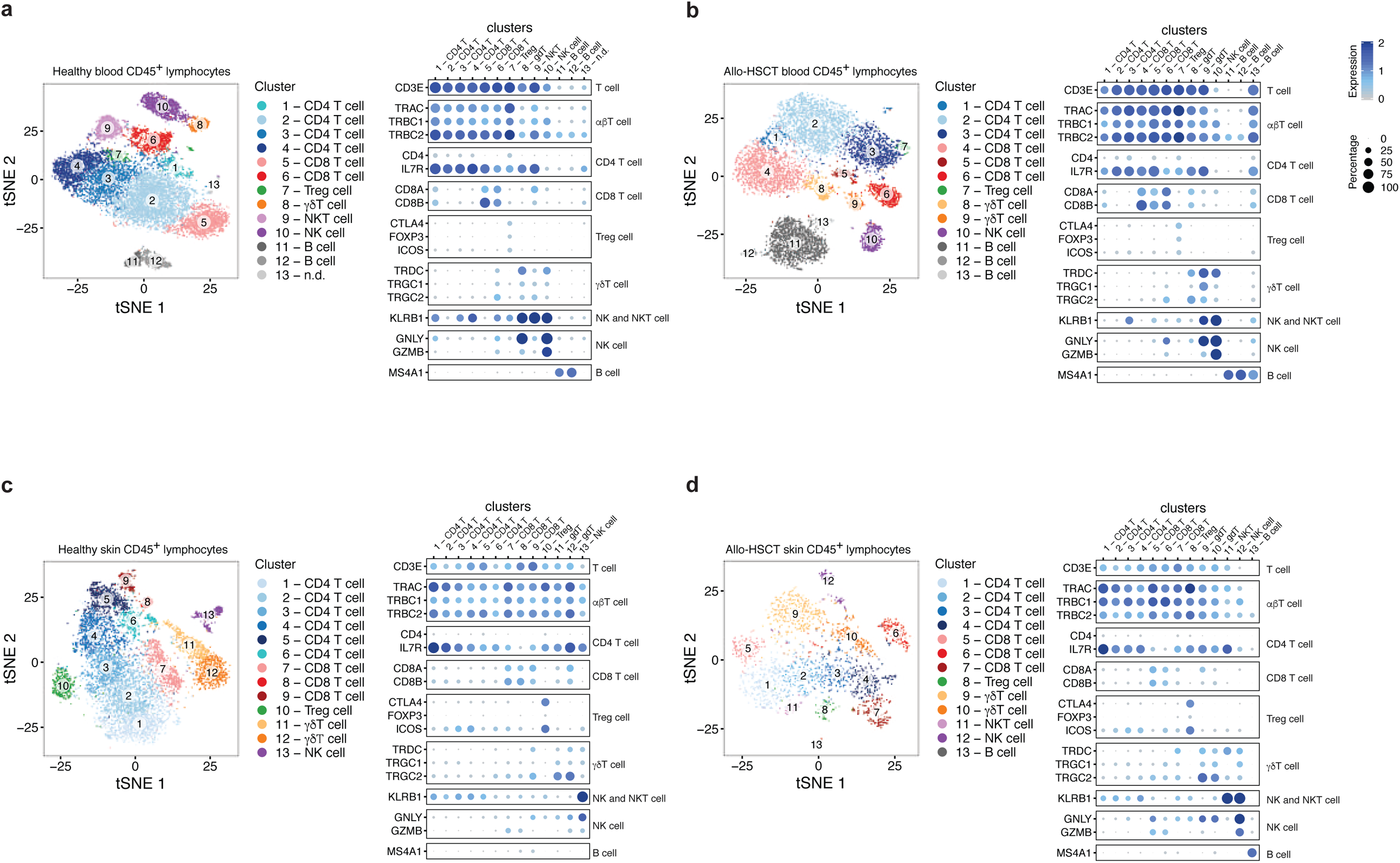
Cell type annotation and clustering of single-cell RNA sequencing data. **(a-d**, left panels) Louvain clusters identified in the blood of the healthy donor depicted in the reduced space calculated by tSNE. **(a-d**, right panels). The cluster annotation in the legend was performed based on the average cluster expression of cell type-specific marker genes. The circle size reflects the percentage of cells expressing the marker genes within the respective cluster.

**Figure S3.**
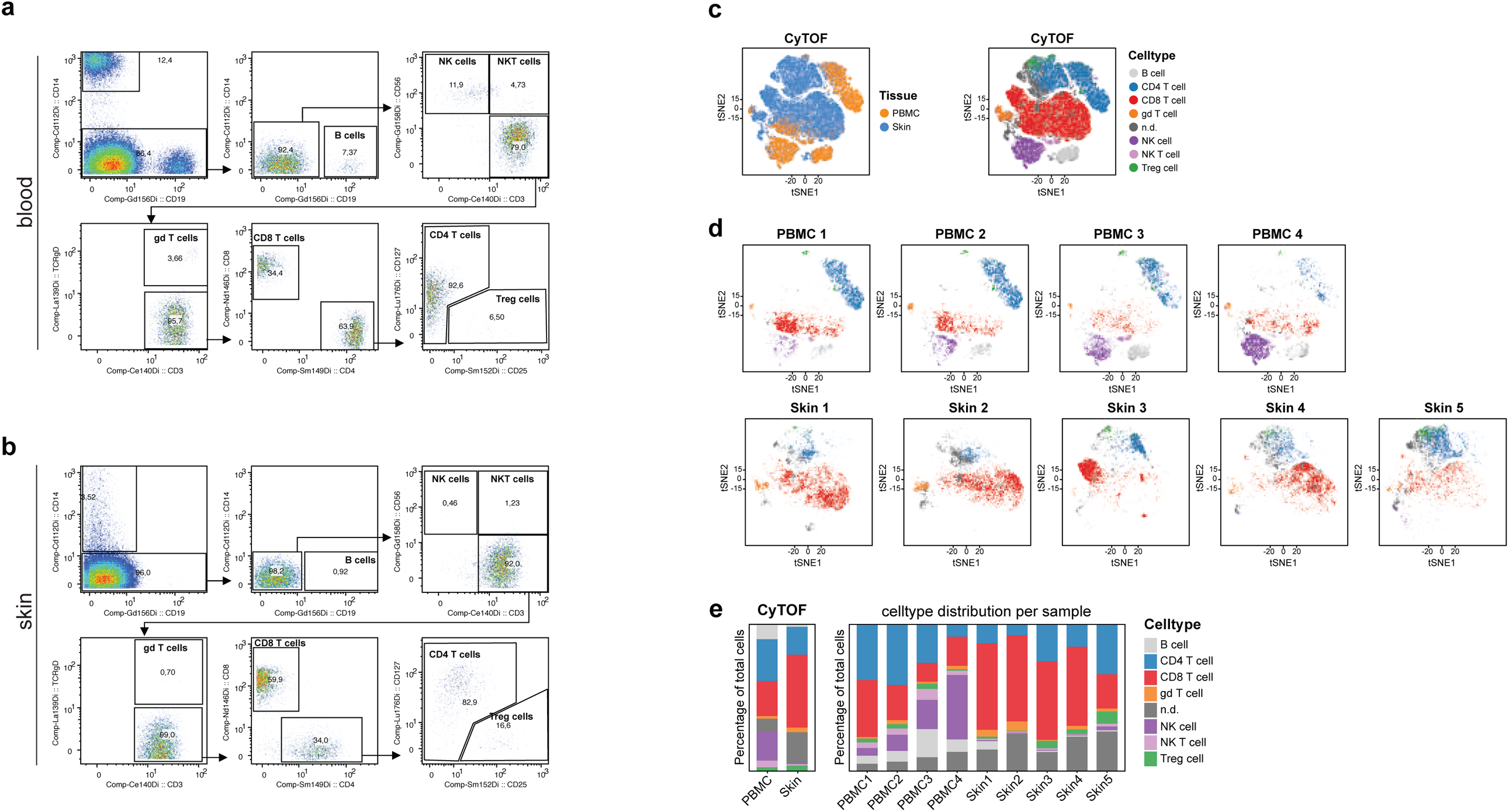
Cell type annotation by CyTOF analysis in healthy skin and blood lymphocytes. **(a-b)** CyTOF gating strategy for blood **(a)** and skin **(b). (c)** Dimensionality reduction by tSNE of normalized protein expression values measured by CyTOF on blood and skin samples from healthy donors (PBMC: n = 4, skin: n = 5). Each donor was downsampled to 11930 total cells. The different colors depict the tissue of origin (left) and the annotated cell type (right) as determined by gating in FlowJo. **(d)** Dimensionality reduction by tSNE of normalized protein expression values measured by CyTOF on blood and skin samples per donor. The different colors depict the annotated cell type as determined by gating in FlowJo. **(e)** Distribution of cell types in skin and blood from healthy donors annotated through manual gating in FlowJo on protein expression values measured by CyTOF. The left plot shows the average distribution over all donors while the right plot depicts the distribution for each donor individually.

**Figure S4.**
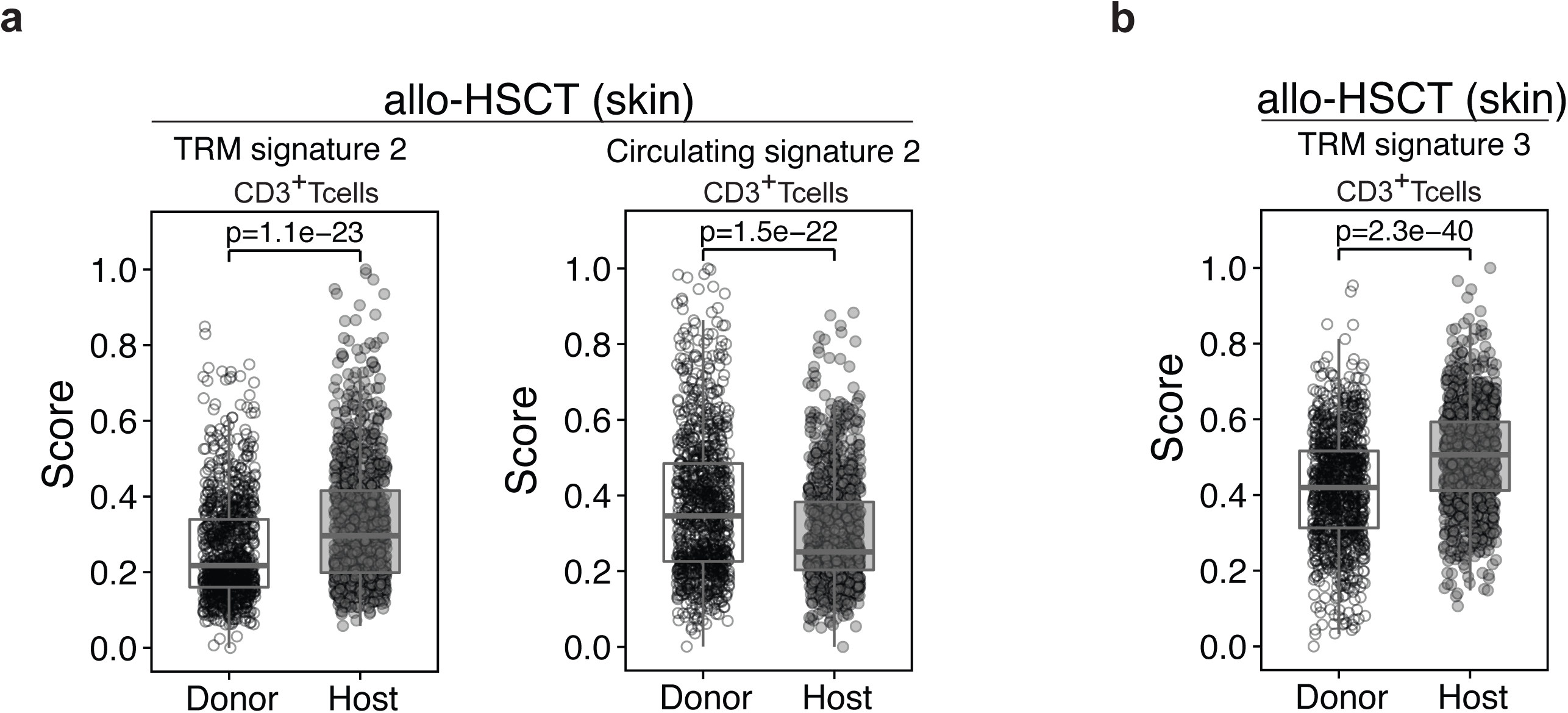
Host lymphocytes in skin and blood after allo-HSCT enrich for different previously established T cell residency signatures. **(a, b)** The CD3^+^ T cells were split between donor and host cells as identified by snpclust and were compared to each other for enrichment of core residency **(a, b)** or core circulating signatures **(a)** as described in Figure 4 using the two-tailed unpaired Student’ t-test. Single-cell gene set expression scores were calculated as the average expression of all genes from the set subtracted by the average expression of random background genes with similar range of expression values. The signatures were generated from publicly available single-cell data sets derived from human cells. **(a)** ^10^, **(b)** ^8^.

**Figure S5.**
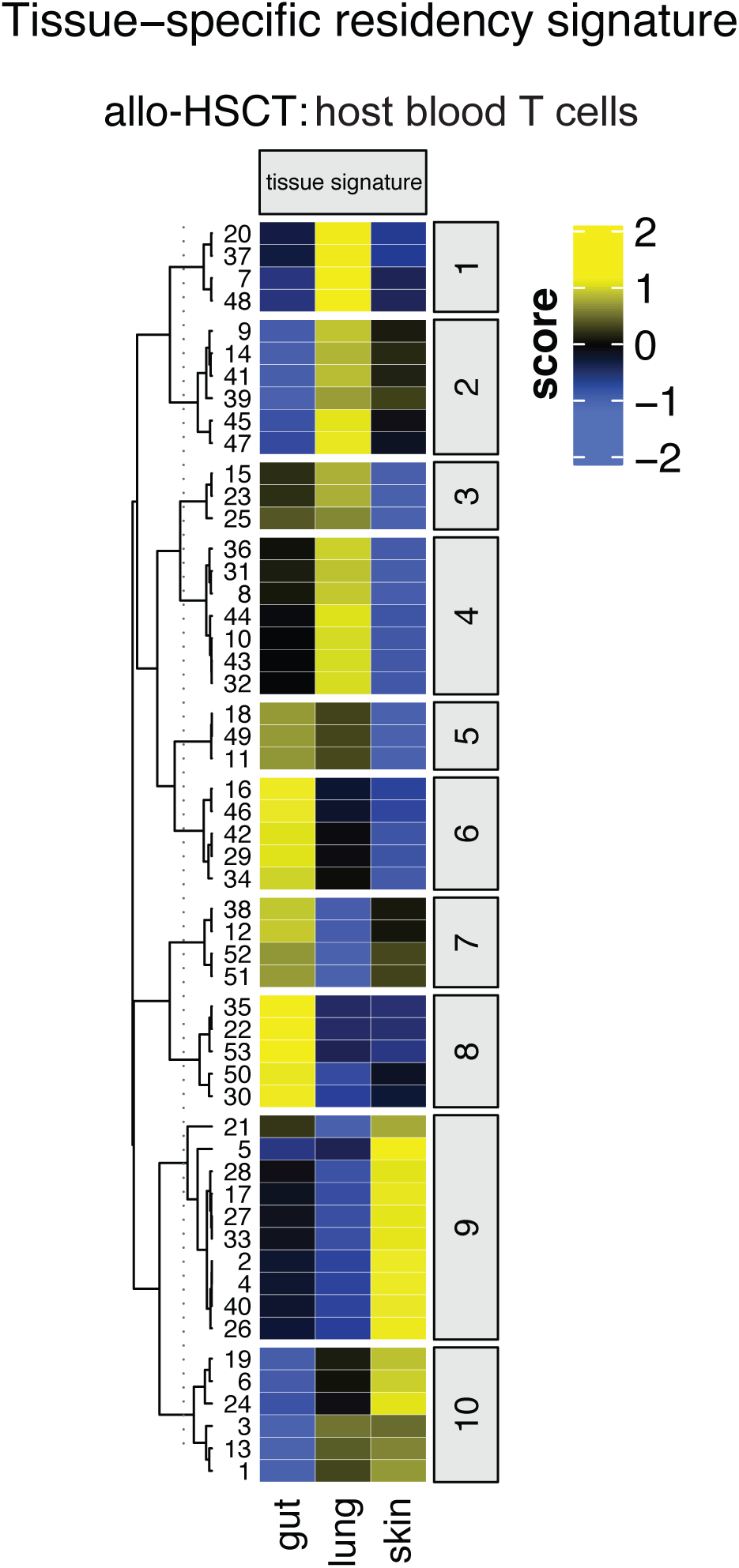
Circulating host blood T cells display a mixed pattern of tissue specific signatures. Single-cell gene set expression scores for tissue-specific residency signatures in each host CD45^+^ lymphocyte detected by SNV and absence of Y-linked genes (53 cells) in the scRNA-Seq from the blood of the allo-HSCT patient. The gene set expression scores were calculated as the average expression of all genes from the set subtracted by the average expression of random background genes with similar range of expression values. The scores were obtained using the AddModuleScore function from Seurat package. Cells were clustered according to their pattern of scores for all gene sets analyzed using k-means clustering (cluster number is indicated on the right side). The numbers on the left indicate the host cell index.

## Acknowledgements

This work was funded by the German Center of Infection Research (to C.E.Z.) and the German Research Foundation (SFB1054, TP10 & SFB1335, TP to C.E.Z). We thank Dr. G. Eckstein for technical support, Julian Reinhard for bioinformatic support, Annika Böttcher for performing the 10xGenomics library preparation and the doctors of the 3. Med. Klinik at Klinikum r. d. Isar in Munich for their support in organizing access to patients.

## Author information

C.E.Z. conceptualized the study, designed the experiments and wrote the manuscript. G.P.A. performed the bioinformatic analysis, P.L. performed STR-PCR and analyzed data, C.F.C. performed experiments, S.M. reanalyzed public CyTOF data.

